# Generalized Normative Modeling: A One-Step Hierarchical Kernel Framework for Multi-Site Brain Charts with Self-Correcting Z-Scores

**DOI:** 10.64898/2026.05.17.725772

**Authors:** Min Li, Ying Wang, Yajun Shen, Lin An, Gangyong Jia, Maria L. Bringas-Vega, Pedro Antonio Valdés-Sosa

## Abstract

Normative modeling expresses individual brain phenotypes as z-scores relative to a population norm, but in multi-site studies batch effects contaminate these z-scores and undermine their biomarker value. Existing approaches either harmonize data before fitting a normative model (ComBat+Normative), letting residual site effects leak into z-scores, or use parametric one-step methods (GAMLSS, HBR) that cannot flexibly model multivariate covariate interactions. We propose **Generalized Normative Modeling (GNM)**, a onestep hierarchical framework that jointly estimates the global trajectory and site-specific effects via NUFFT-accelerated kernel regression with GCV bandwidth selection. Because the z-score is the ratio of batch-corrected residual to batch-corrected scale, residual site variance cancels algebraically — a property we term *self-correction*. On ABIDE I cortical thickness (387 HC, 11 sites, 68 ROIs) and HarMNqEEG log-power spectra (1,564 subjects, 14 sites, 18 channels × 235 frequency bins), GNM produced the most site-invariant z-scores and best age-signal preservation among four methods. This work provides an open-source MATLAB toolbox with a declarative formula interface (https://github.com/LMNonlinear/Generalized-Normative-Modeling), enabling reliable individual-level inference in pooled multi-site cohorts and advancing the use of normative deviations as clinical biomarkers in precision psychiatry and neurology.

## 1. Introduction

In medicine and neuroscience, normative modeling has emerged as a framework that goes beyond group-level comparisons to capture individual neurobiological “fingerprints” (Marquand et al., 2016, 2019). The core idea is straight-forward: given a brain-derived descriptive parameter (DP) — such as cortical thickness, spectral power, or connectivity — and a set of clinically relevant parameters (RCPs) such as age, the normative model estimates how the DP varies across the healthy population. Each individual is then assigned a z-score that quantifies their deviation from the population norm. This individual-level characterization has proven valuable for studying the biological heterogeneity of psychiatric and neurological conditions, where patients deviate from the norm in different directions and at different brain locations — patterns that cancel out in conventional group analyses (Wolfers et al., 2018; Zabihi et al., 2019).

The most prominent application of normative modeling is the construction of developmental brain charts — analogous to pediatric growth charts for height and weight — that describe how brain phenotypes change with age across the lifespan (Bethlehem et al., 2022). By establishing these normative trajectories from large samples of healthy subjects, researchers can conceptualize mental disorders as deviations from the expected developmental benchmark. This approach has been applied to detect white matter injury in infants, characterize heterogeneity in schizophrenia and bipolar disorder, and construct brain-predicted age as a biomarker for neurodegeneration and mortality (Cole et al., 2017; Cole and Franke, 2017). Combined with genetic and clinical data, normative z-scores provide a quantitative bridge between population-level norms and individual-level pathology.

The procedure of normative modeling involves four steps: (1) collect data and extract DPs that capture physiological or structural features; (2) estimate the normative function — the “norms” — that describes how DPs change with RCPs, including both the central tendency and the dispersion; (3) compute individual z-scores on held-out data to validate the model; and (4) use the z-scores for downstream statistical analysis, prediction, or classification. The second step — estimating the norms — is the most critical, and it is here that several statistical challenges arise.

### Nonlinearity

The relationship between brain measures and age is typically nonlinear: cortical thickness increases rapidly in childhood, peaks in adolescence, and declines through adulthood (Bethlehem et al., 2022). Simple linear regression is inadequate for capturing these trajectories, biasing z-scores at both ends of the age range, and motivating the use of nonparametric smoothers such as P-splines, kernels, or Gaussian processes (Rigby and Stasinopoulos, 2005; Marquand et al., 2019).

### Heteroscedasticity

Regression models typically assume equal variance across samples. However, inter-individual variability in brain measures changes with age — younger and older subjects tend to show greater variability than middle-aged adults (Bethlehem et al., 2022; Marquand et al., 2019). When the data violate the homoscedasticity assumption, z-scores computed with a constant variance estimate are systematically inflated or deflated depending on the individual’s position along the covariate axis. Joint modeling of location and scale as functions of the covariates, as in GAMLSS (Rigby and Stasinopoulos, 2005; Dinga et al., 2021), is required to restore calibration.

### Non-Gaussian distributions

The DPs are not always Gaussian. It has been shown that Gaussianity of the residuals is needed to avoid false positives in z-score-based inference (Fraza et al., 2021; de Boer et al., 2024). For non-Gaussian DPs, link functions or distributional transformations (Box–Cox, sinh-arcsinh) are needed to map the data to a scale where the location-scale z-score framework is valid (Rigby and Stasinopoulos, 2005; Fraza et al., 2021; de Boer et al., 2024).

### Batch effects

In the era of open science, multi-site data pooling has become essential for achieving the sample sizes and demographic diversity that normative models require. Consortia such as ABIDE (Di Martino et al., 2014), ENIGMA (Thompson et al., 2014), the UK Biobank (Miller et al., 2016), and HarMN-qEEG (Li et al., 2022) aggregate thousands of recordings from dozens of sites. However, pooling introduces non-biological sources of variability — differences in recording equipment, acquisition protocols, and preprocessing pipelines — collectively termed “batch effects.” These effects can account for 20–60% of the variance in derived brain measures (Fortin et al., 2018; Pomponio et al., 2020), often exceeding the biological signals of interest. For multi-site normative modeling, the model must be extended from a fixed-effect structure to an additive mixed-effect model that incorporates batch as a random effect alongside the biological fixed effects.

### Fast multivariate regression

As datasets grow to tens of thousands of subjects with high-dimensional DPs, traditional regression algorithms with 𝒪(*N* ^2^) complexity become computationally prohibitive. Fast algorithms — such as those based on the Fast Fourier Transform — are needed to serve as the computational backbone of the normative modeling pipeline.

These challenges have motivated several recent methodological advances. The prevailing approach for batch effect removal is ComBat (Johnson et al., 2007; Fortin et al., 2017, 2018), which models additive and multiplicative site effects using empirical Bayes shrinkage. ComBat effectively removes site-driven variance while preserving specified biological covariates, but it produces only harmonized data values — not normative z-scores. Obtaining z-scores requires a separate normative modeling step, creating a two-step pipeline (harmonize, then model) in which residual batch effects from the first step propagate into the second.

To achieve tighter integration, Hierarchical Bayesian Regression (HBR) (Kia et al., 2022) treats site as a random effect within the normative model, allowing batch and normative parameters to be estimated simultaneously. Fraza et al. (2021) extended this line with warped Bayesian linear regression for non-Gaussian distributions, and de Boer et al. (2024) introduced the sinh-arcsinh (SHASH) likelihood to model all four distribution parameters (location, scale, skewness, kurtosis) as functions of covariates and site. From a different perspective, GAMLSS (Rigby and Stasinopoulos, 2005; Dinga et al., 2021) extends generalized additive models to jointly estimate location, scale, and shape parameters, with site as a fixed or random effect; the Brain Charts project (Bethlehem et al., 2022) applied GAMLSS at unprecedented scale (∼120,000 MRI scans). On the deep learning side, methods such as DeepComBat (Hu et al., 2024), adversarial unlearning (Dinsdale et al., 2021), and conditional VAEs (Moyer et al., 2020) have been proposed, but these approaches typically sacrifice the explicit distributional framework needed for calibrated z-score computation.

Despite this progress, a gap remains. All these methods — HBR, GAMLSS, ComBat, and their extensions — can be understood as special cases of a single unified equation for harmonized normative modeling, recently formalized as a location–scale framework (Li et al., 2026a,b):

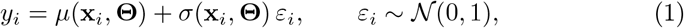

where *µ*(**x, Θ**) and *σ*(**x, Θ**) are the location and scale functions that depend on both biological covariates **x** and nuisance parameters **Θ** (including batch effects), and the z-score is 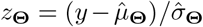. ComBat restricts *µ* to a linear function of covariates with constant batch shifts; GAMLSS allows nonparametric additive terms but typically uses linear batch effects; HBR places Bayesian priors on the batch parameters but relies on parametric basis functions. No existing method simultaneously provides: (i) fully nonparametric estimation of both *µ* and *σ* as multivariate functions of covariates, (ii) flexible batch effect modeling (constant or covariate-dependent), and (iii) computational scalability to large datasets — all within a single unified framework.

In this work, we propose **Generalized Normative Modeling (GNM)**, a one-step framework that instantiates this unified equation using nonparametric kernel regression with a hierarchical batch structure (fig. 1). The framework has three defining features:

**Figure 1.**
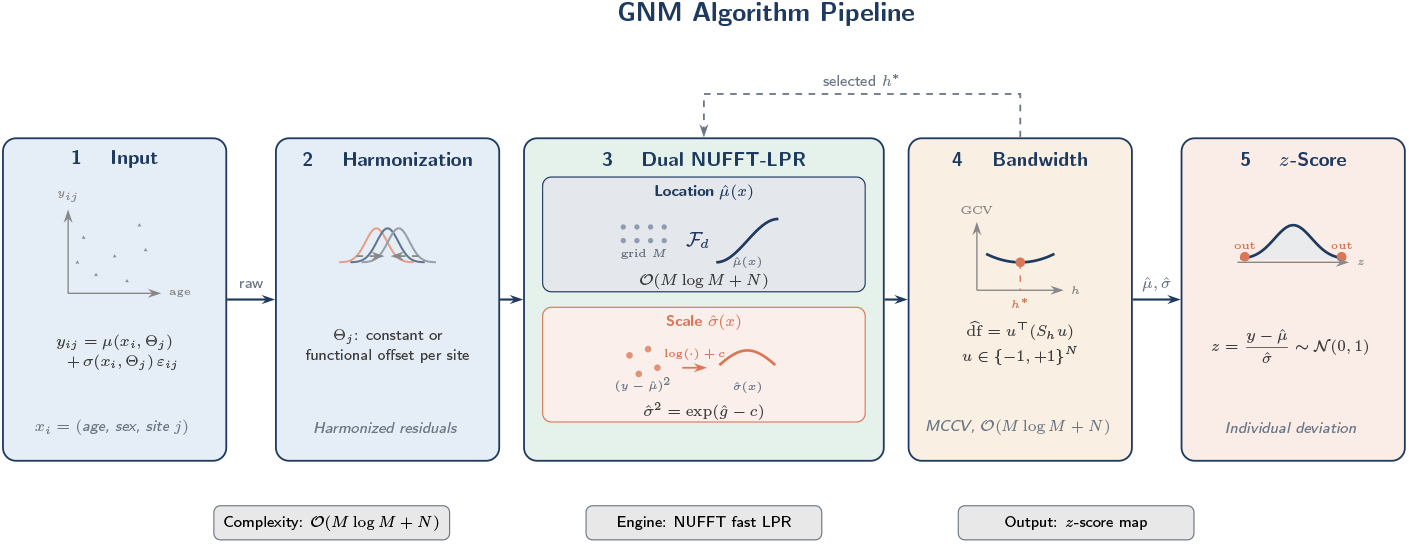
GNM algorithm pipeline. Five stages instantiating the location–scale form *y*_ij_ = *µ*(*x*_i_, Θ_j_) + *σ*(*x*_i_, Θ_j_) *ε*_ij_, *ε*_ij_ ∼ 𝒩 (0, 1). **(1) Input**: multi-site brain features *y*_ij_ with covariates *x*_i_ = (age, sex, site *j*). **(2) Harmonization**: site batch effects Θ_j_ absorbed as constant or covariate-functional offsets. **(3) Dual NUFFT-LPR**: 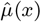 via fast local polynomial regression on an equispaced grid (ℱ_d_, 𝒪(*M* log *M* + *N*)); 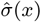 via a second LPR pass on the log-residual *g*(*x*) = log *σ*^2^(*x*) + *c* with back-transform 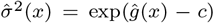. **(4) Bandwidth selection**: GCV minimized via Girard’s stochastic trace estimator 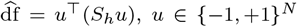; optimal *h*^⋆^ fed back to stage 3 (dashed loop). **(5)** *z***-score & deviation**: 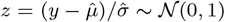 for healthy reference subjects; tail dots mark candidate outliers. End-to-end complexity 𝒪(*M* log *M* + *N*).

1. **Nonparametric kernel regression**. GNM estimates *µ*(**x**) and *σ*(**x**) using local polynomial regression accelerated by the Non-Uniform Fast Fourier Transform (NUFFT), reducing computational complexity from 𝒪 (*N* ^2^) to 𝒪 (*N* log *N*). The smoothing bandwidth is selected automatically via generalized cross-validation (GCV), eliminating manual specification of basis functions, knot locations, or spline orders. Crucially, the kernel smoother natively supports *multivariate* covariates (e.g., joint frequency × age surfaces), capturing interaction effects that per-bin or additive models cannot represent. Existing methods — ComBat, GAMLSS, HBR — process each frequency bin or ROI independently, discarding cross-covariate structure; GNM models the full *d*-dimensional covariate surface in a single pass.
2. **Hierarchical batch model**. Site effects are modeled as a second level within the same estimation, operating on the z-score residuals of the global normative model. Two configurations are supported: a *constant* batch model (subsumes ComBat) and a *functional* batch model where site effects vary smoothly with covariates (generalizes GAMLSS). Because both levels are estimated jointly, the normative trajectory is informed by the batch structure and vice versa, avoiding the error propagation inherent to two-step pipelines.
3. **Self-correcting z-scores**. The z-score is computed as the ratio of the batch-corrected residual to the batch-corrected standard deviation. Because both numerator and denominator incorporate the same batch estimates, residual site-related variance cancels algebraically — the property we term *self-correction*. This means that even when data-level harmonization is imperfect, the z-scores can still be site-invariant by construction.

We validate GNM on two multi-site datasets spanning different neuroimaging modalities. The first is the ABIDE I cortical thickness dataset (387 healthy controls, 11 sites, 68 ROIs), a standard benchmark in the harmonization literature. The second is the HarMNqEEG log-power spectral dataset (1,564 subjects, 14 sites, 18 EEG channels, 235 frequency bins), which presents a higher-dimensional challenge requiring 2-D kernel regression over the joint frequency × age surface. On both datasets, we compare GNM against ComBat + Normative model, GAMLSS, and Hierarchical Bayesian Regression (HBR), evaluating performance across three dimensions: (i) batch effect removal, (ii) biological signal preservation, and (iii) z-score calibration and cross-site comparability.

## 2. Methods

### 2.1. The unified equation of harmonized normative modeling

Let *y*_*ij*_ denote the observation for subject *i* at site *j*, and let **x**_*i*_ denote the vector of clinically relevant parameters (e.g., age, or jointly frequency and age). The observation can be decomposed as:

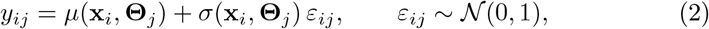

where *µ*(**x, Θ**) is the location function (normative mean), *σ*(**x, Θ**) is the scale function (normative standard deviation), and **Θ**_*j*_ denotes the nuisance parameters associated with site *j*. The normative z-score is:

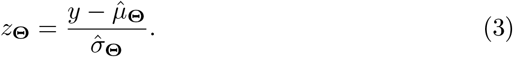

If *µ* and *σ* are estimated accurately, *z* ∼ 𝒩 (0, 1) for healthy subjects, and the z-score is independent of both the biological covariates **x** and the batch identity *j*. Equation (2) provides a unified framework that subsumes existing harmonization and normative modeling methods as special cases, differing only in the functional forms assumed for *µ* and *σ* and in how the batch parameters **Θ** are estimated (section 2.7).

### 2.2. The generalized harmonized normative model

For multi-center data, the factors influencing a DP extend beyond biological covariates to include nuisance grouping variables such as sex and batch. Let **x**_*i*_ denote the user-specified biological covariates and **r**_*i*_ = (*r*_*i*1_, *r*_*i*2_, …) denote grouping factors treated as random effects. The generalized harmonized normative model is written as:

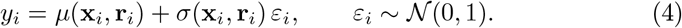

To separate the biological fixed effects from the random effects, the location and scale functions are decomposed as:

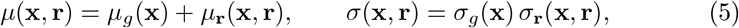

where *µ*_*g*_, *σ*_*g*_ are the global (population-level) normative functions of the biological covariates, while *µ*_**r**_, *σ*_**r**_ capture the additive and multiplicative random effects. Substituting into eq. (4):

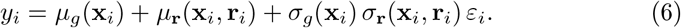

Focusing on batch as the primary random effect and writing *j* for the batch identity:

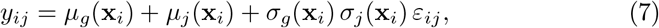

where *µ*_*j*_(**x**) is the batch-specific additive shift on the mean and *σ*_*j*_(**x**) is the batch-specific multiplicative scaling on the variance.

### 2.3. Hierarchical solving strategy

#### 2.3.1. Rescaling for hierarchical z-score computation

To enable layered z-score computation, we rescale the batch-level parameters by the global standard deviation:

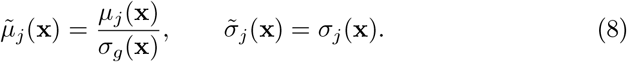

Substituting into eq. (7) and factoring out *σ*_*g*_(**x**_*i*_):

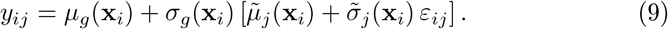

This form separates the model into two nested levels: the outer level describes the global normative trajectory *µ*_*g*_, *σ*_*g*_, while the inner bracket describes the batch-specific deviation 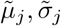 on a standardized scale.

#### 2.3.2. Decomposition reveals existing methods as special cases

The decomposition in eq. (9) implies:

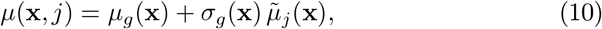

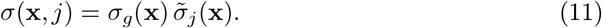

This reveals that the unified equation contains existing methods as special cases:

- When *µ*_*g*_ is *linear* in covariates, *σ*_*g*_ is *constant*, and 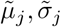 are *constants*, the model reduces to **ComBat**.
- When 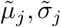 are *linear* in covariates, the model reduces to **GAMLSS** with site as a random effect.
- In **GNM**, both fixed-effect and random-effect functions are estimated *non-parametrically* via kernel regression.

The automatic GCV bandwidth selection (section 2.5) implicitly performs model comparison: when the data support only a constant batch effect, the GCV-selected bandwidth becomes maximal (yielding a flat function ≡ constant); when the data support covariate-dependent batch effects, the bandwidth becomes smaller, recovering a smooth function. Thus GNM subsumes the functional forms of both ComBat and GAMLSS as limiting cases within a single estimation framework.

#### 2.3.3. Step-by-step solving procedure

The hierarchical model (eq. (9)) is solved in four steps (fig. 2).

**Figure 2.**
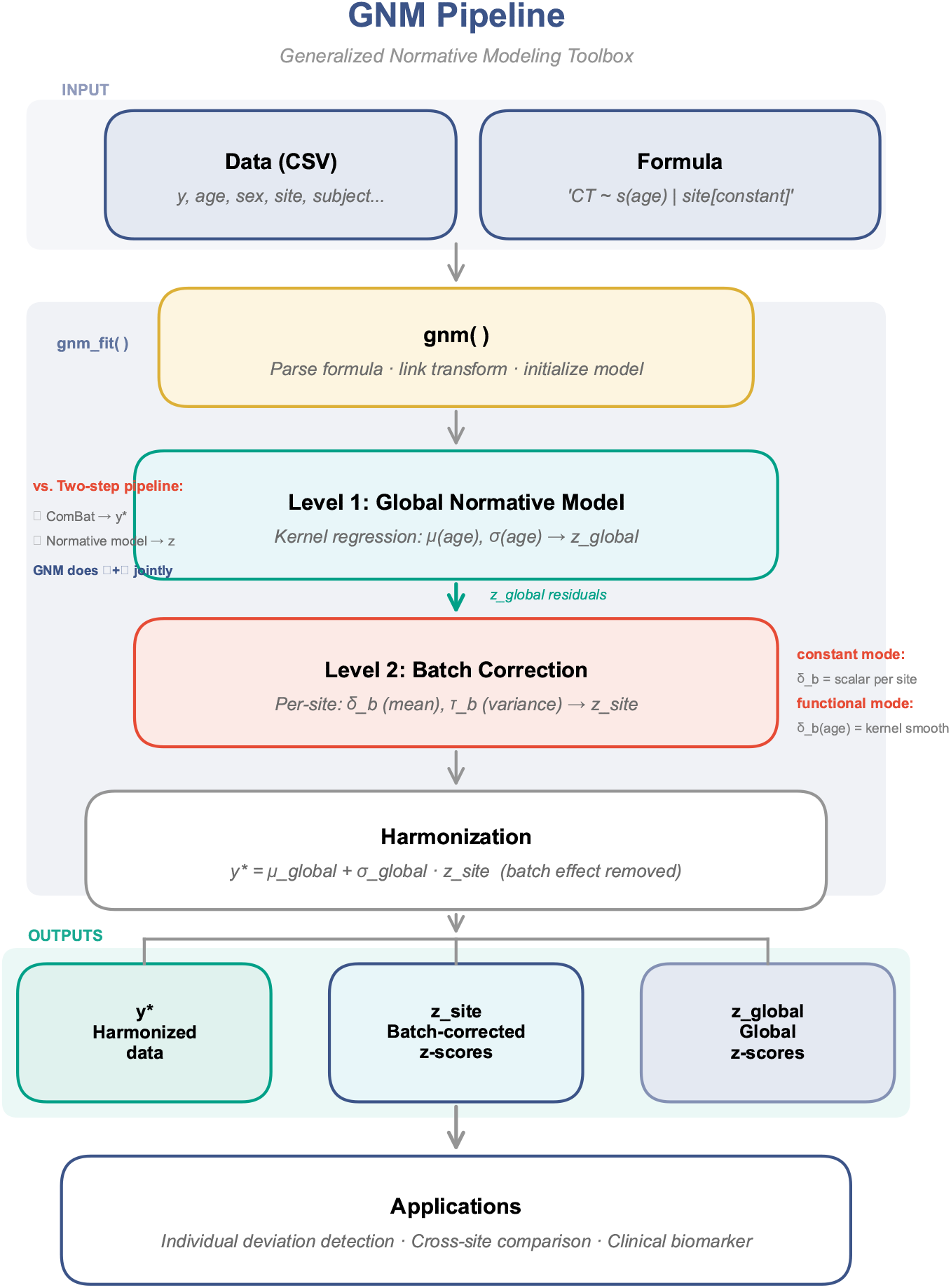
GNM algorithm schematic. The pipeline processes multi-site data jointly: the global normative trajectory and site-specific batch effects are estimated in a single hierarchical kernel-regression step, yielding self-correcting z-scores without an upstream harmonization stage.

##### Step 1

Pooling all subjects, fit the global mean and standard deviation,

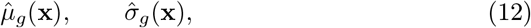

both estimated by NUFFT-accelerated kernel regression (section 2.4): 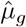 smooths *y* over **x**, and 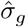 smooths the squared residuals 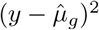 over **x**.

##### Step 2

Compute the global z-scores:

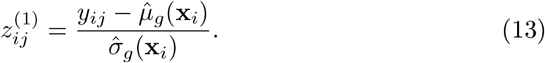

##### Step 3

For each batch *j*, estimate the batch-specific rescaled mean 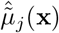 and standard deviation 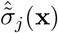 from 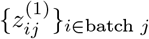. The batch-corrected z-score is:

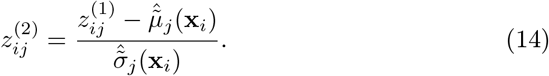

##### Step 4

Reconstruct the harmonized DP by substituting 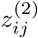 back through the global model:

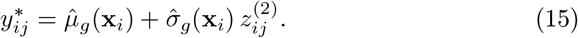

The harmonized normative trajectory (brain chart) is obtained by smoothing *y*^∗^ over the covariates.

*Batch model configurations*.. The functional form of 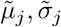 in Step 3 is specified through the formula interface:

- *Constant batch* (`batch_var[constant]`): 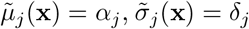— scalar constants per batch.
- *Functional batch* (`batch_var[functional(x1)]`): 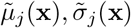 are smooth functions of specified covariates, estimated by kernel regression within each batch.

##### Self-correcting property

Because 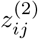 is the ratio of two batch-corrected quantities (eq. (14)), any residual site effect in 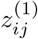 that is captured consistently by both 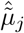 and 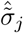 cancels algebraically. This means that even when data-level harmonization is imperfect (i.e., *y*^∗^ retains some site structure), the z-scores can still be perfectly site-invariant.

### 2.4. NUFFT-accelerated kernel regression

All smooth functions in the hierarchical model — 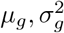 at Level 1 and 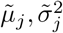 at Level 2 — are estimated by *local polynomial regression* (LPR) of order *l*, accelerated by the Non-Uniform Fast Fourier Transform (NUFFT) (Fan and Gijbels, 1996; Wang et al., 2022). The implementation builds on the FKreg toolbox (Wang et al., 2022), which has been previously applied to model nonlinear resonance effects in human cortical oscillations (Wang et al., 2026).

Given *N* observations 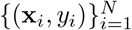 with **x**_*i*_ ∈ ℝ^*d*^, the LPR estimator at a target point **x** minimizes

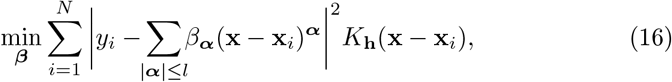

where 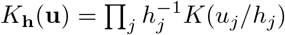 is a product Epanechnikov kernel. The closed-form solution is

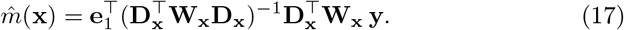

The entries of 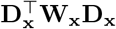 and 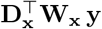 are kernel-weighted polynomial moments — discrete convolution sums:

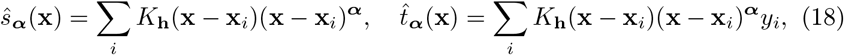

so that the LPR estimate at grid point **g**_*m*_ is 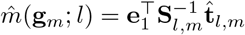.

The convolution structure of eq. (18) enables NUFFT acceleration: direct evaluation costs 𝒪(*NM*), but GNM computes the moments in the frequency domain at 𝒪(*N* + *M*^*d*^ log *M*). Define the type-1 nonuniform Fourier transforms 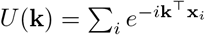 and 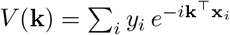. By the convolution theorem, the moments in the frequency domain become 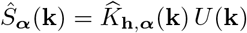 and 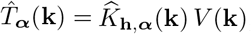, recovered to the grid by inverse FFT. The type-1 transforms are computed via fast Gaussian gridding. After computing 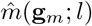 on the regular grid, estimates at the original sample locations are recovered by spline interpolation.

### 2.5. Bandwidth selection via GCV

The smoothing bandwidth **h** is selected using generalized cross-validation:

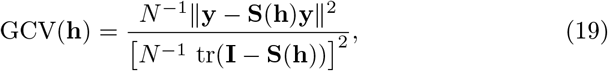

where **S**(**h**) is the smoother matrix and df(**h**) = tr(**S**(**h**)).

Forming **S**(**h**) explicitly costs 𝒪(*N* ^2^) memory. GNM estimates the trace via Hutchinson’s stochastic estimator using *n*_*p*_ random probe vectors **z**_*j*_ ∼ 𝒩 (**0, I**):

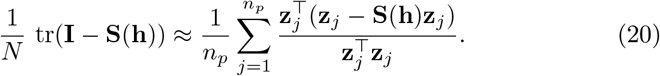

Each **S**(**h**)**z**_*j*_ is computed by the same NUFFT pipeline, so the GCV cost is comparable to a single fit; the default is *n*_*p*_ = 10. A candidate bandwidth grid is specified per covariate dimension, and the optimal bandwidth is selected using the *one-standard-error (1-SE) rule*: among bandwidths whose GCV score falls within one SE of the minimum, the smoothest (largest bandwidth) is chosen, which regularizes toward smoother estimates and reduces overfitting.

#### Variance estimation

The conditional variance is estimated via log-space LPR with multiplicative bias correction (Fan and Yao, 1998): 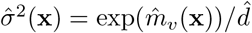, where 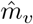 is the LPR fit of 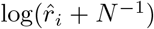 on the squared residuals 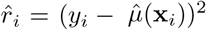, and 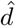 is a multiplicative bias-correction factor.

### 2.6. Link functions, outlier detection, and prediction

#### Link functions

When the DP is not approximately Gaussian on the natural scale, a monotonic link function *g*(·) transforms the response before fitting (identity, log, logit, sqrt, probit). The pipeline applies *g*(*y*) before kernel regression and *g*^−1^ to harmonized values *y*^∗^ afterward; z-scores remain on the standard normal scale by construction (fig. 3).

**Figure 3.**
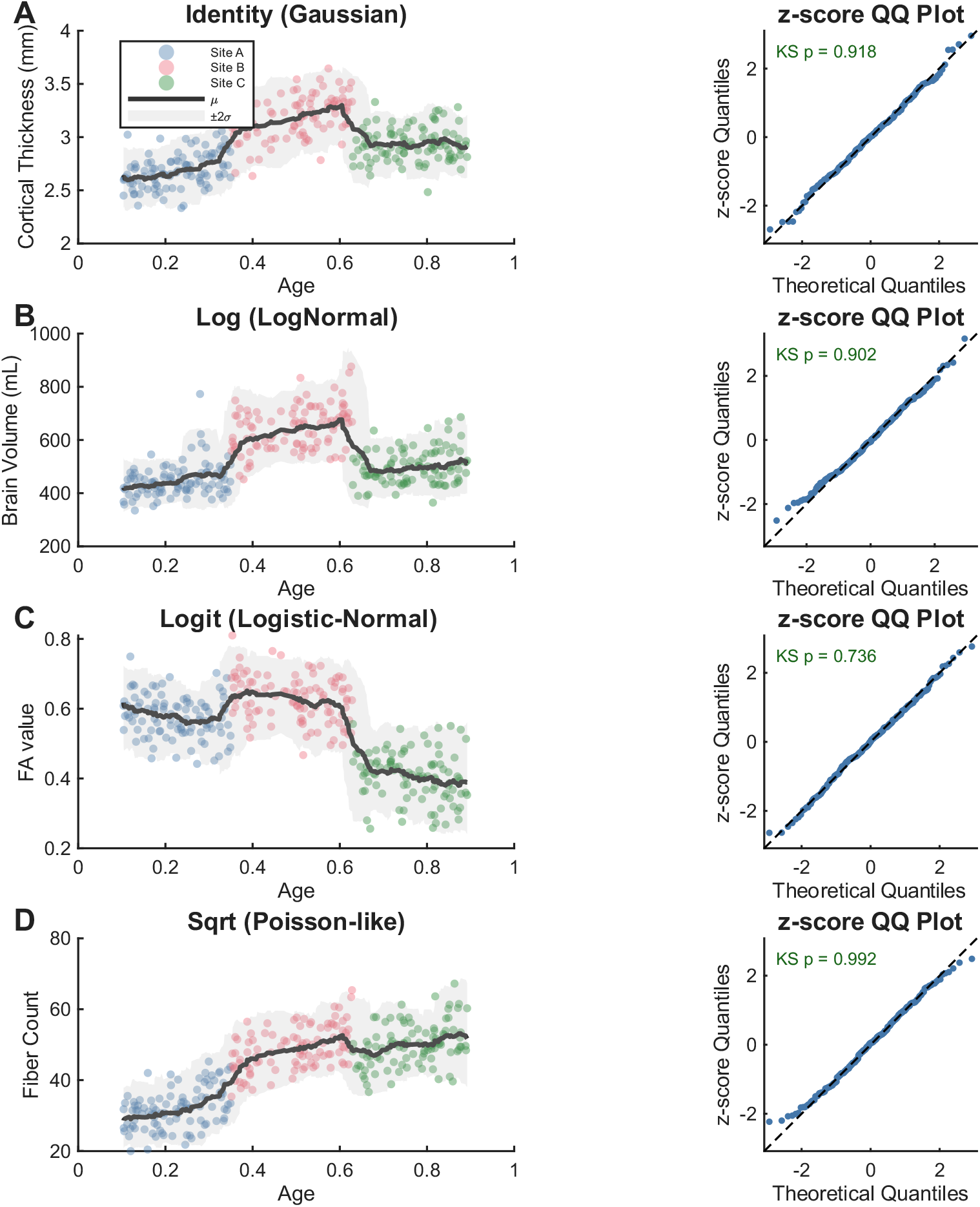
Link functions map non-Gaussian response variables to a scale on which the location-scale z-score framework is valid. Shown are identity (Gaussian), log (positive-only), and logit (bounded) links applied to simulated responses.

#### Robust outlier detection

An optional preprocessing step identifies multivariate outliers using the Minimum Covariance Determinant (MCD) estimator (Rousseeuw and Van Driessen, 1999) operating on the *global z-scores* 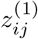 rather than raw observations *y*_*ij*_. This decouples outlier detection from the covariate trajectory and the batch structure. Multivariate z-score profiles are projected to two dimensions via t-SNE, and observations exceeding 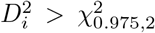 are flagged (fig. 4). This step is controlled by `clean_individual_outlier` (default: off).

**Figure 4.**
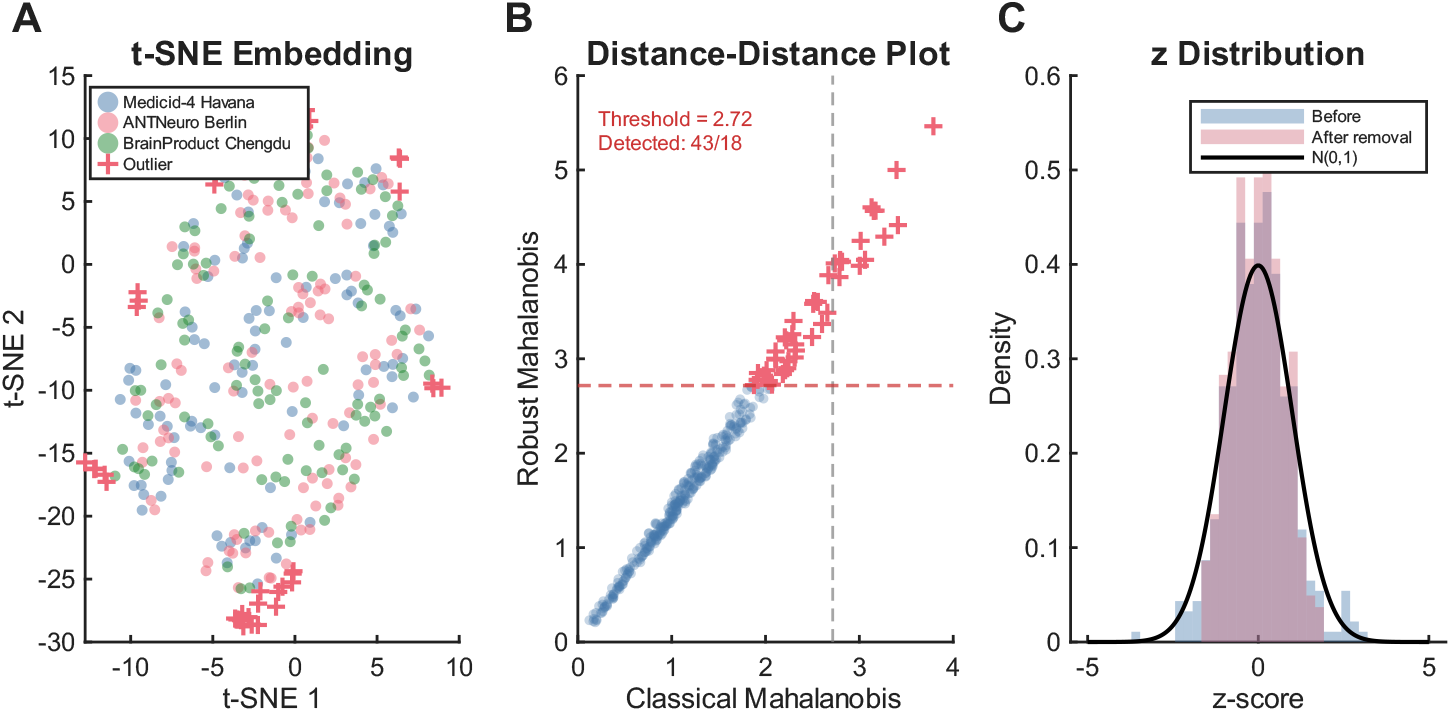
Robust outlier detection via MCD-based Mahalanobis distance in the z-score space. Contaminated observations are down-weighted prior to batch-effect estimation, preventing inflation of site-specific scale parameters.

#### Prediction on new data

A trained GNM model can be applied to new (out-of-sample) data without retraining. For an observation *y*_*i*,new_ from a known batch *j* with covariates **x**_*i*,new_, the stored global estimates are evaluated at **x**_*i*,new_ via grid interpolation, then the stored batch estimates yield *z*^(2)^ (eq. (14)) and *y*^∗^ (eq. (15)). For new data from a batch *j*^′^ not in training, the user can specify a reference batch *j*_ref_ via the `reRefBatch` mechanism: 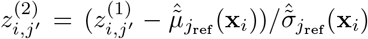.

### 2.7. Relationship to existing methods

#### 2.7.1. ComBat + Normative model (two-step)

ComBat (Johnson et al., 2007; Fortin et al., 2017) models

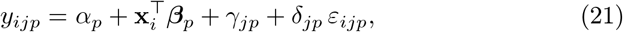

with *γ*_*jp*_, *δ*_*jp*_ as *constant* batch effects estimated by empirical Bayes shrinkage. ComBat produces harmonized data *y*^∗^ but does *not* produce z-scores directly. A separate normative model fits 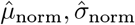 to *y*^∗^, and the batch-adjusted z-score is 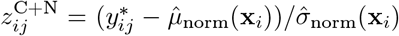. Because the second-step model has no access to the original batch structure, any residual batch effect in *y*^∗^ propagates directly into *z*^C+N^.

#### 2.7.2. GAMLSS (one-step, parametric)

GAMLSS (Rigby and Stasinopoulos, 2005; Dinga et al., 2021) models each distribution parameter as smooth additive functions of covariates plus site effects:

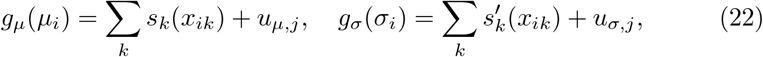

with P-spline smoothers *s*_*k*_ and site terms *u*_*j*_. GAMLSS provides one-step z-scores. However, the additive structure (e.g., pb(freq)+pb(age)) cannot capture covariate interactions without explicit tensor product terms, and site effects are typically restricted to linear terms.

#### 2.7.3. Summary

Table 1 contrasts the four methods along the parameterization axes of eq. (2).

**Table 1.** Comparison of harmonized normative modeling methods.

| Method | $\mu$ fixed | $\mu$ batch | $\sigma$ fixed | $\sigma$ batch | z-score | Estimation |
| --- | --- | --- | --- | --- | --- | --- |
| ComBat+ Norm | linear | constant | — | constant | two-step | EB + separate rate |
| GAMLSS | additive P-spline | linear | additive P-spline | linear | one-step | Penalized likelihood |
| HBR | parametric basis | hierarchical | — | hierarchical | one-step | MCMC |
| **GNM** | **nonparametric** | **const./functional** | **nonparametric** | **const./functional** | **one-step** | **GCV + NUFFT** |

The fundamental distinction is between *one-step* methods (GNM, GAMLSS, HBR), which produce batch-adjusted z-scores directly from the joint model, and the *two-step* pipeline (ComBat+Norm), which harmonizes data first and then fits a separate normative model. Among one-step methods, GNM is the most flexible: nonparametric kernel regression for both fixed and random effects, native multivariate covariate support, and automatic batch-model complexity selection via GCV. All four methods are empirically compared on ABIDE I (section 3.2.1), and three (excluding HBR) on HarMNqEEG (section 3.3).

### 2.8. Software and formula interface

GNM is implemented as an open-source MATLAB toolbox (GNM-ToolBox).

The complete pipeline is driven by a formula interface:

~~~
% Constant batch (subsumes ComBat)
mnhs = gnm(‘data.csv’, ‘CT ∼ s(age) | site[constant]’);
% Functional batch (generalizes GAMLSS)
mnhs = gnm(‘data.csv’, ‘logpower ∼ s(freq, age) | site[functional(age)]’);
mnhs = gnm_fit(mnhs); % train
T_new = gnm_predict(mnhs, ‘new_data.csv’); % predict
~~~

The formula parser extracts response variables (left of ∼), smooth terms `s`(…) defining covariates, the batch variable (after |), and the batch model type (`constant` or `functional`). The pipeline proceeds: parse formula → initialize model → optional link transform → Level 1 global kernel regression → Level 2 batch correction → harmonization → inverse link transform → output (*y*^∗^, *z*_global_, *z*_site_); see fig. 5.

**Figure 5.**
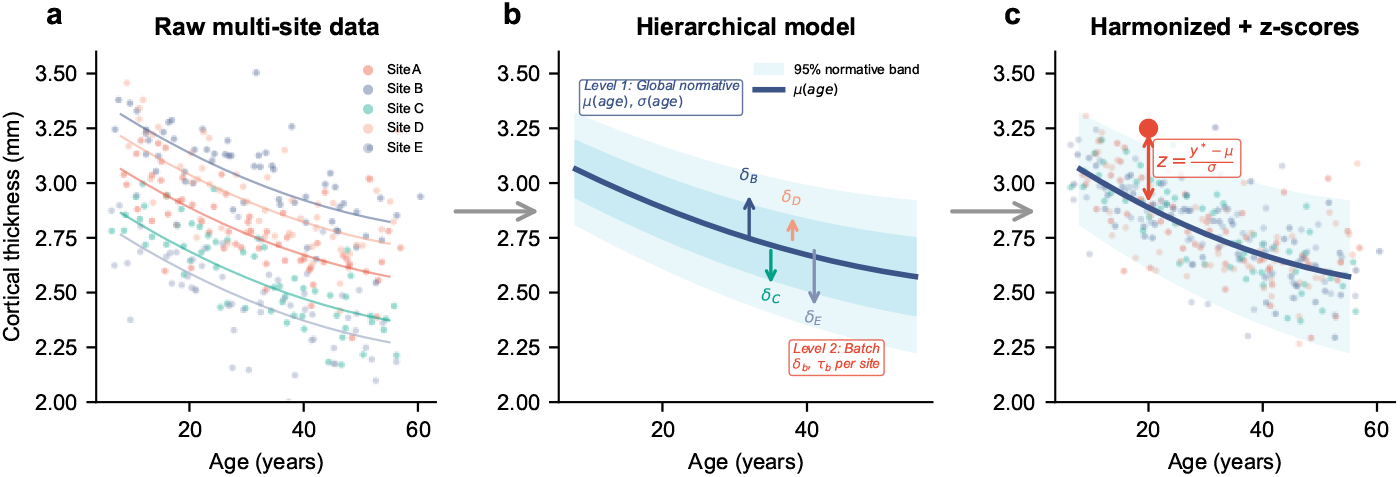
Hierarchical model decomposition. (a) Raw multi-site data with visible batch offsets. (b) The two-level model: Level 1 estimates the global normative band *µ*(*age*), *σ*(*age*); Level 2 estimates per-site batch shifts *δ*_b_ and scales *τ*_b_ on the standardized residuals. (c) Harmonized data with batch effects removed and individual z-scores computed.

### 2.9. Datasets

Two multi-site neuroimaging datasets spanning different modalities, spatial scales, and dimensionalities were used to evaluate GNM.

#### ABIDE I (structural MRI)

Cortical thickness (CT) measurements were obtained from the Autism Brain Imaging Data Exchange I (Di Martino et al., 2014), parcellated with FreeSurfer using the Desikan-Killiany atlas (68 ROIs, 34 per hemisphere). From the initial pool of 528 healthy controls (HC) across 20 sites, sites with fewer than 20 subjects were excluded, yielding a final sample of 387 HC across 11 sites (age range: 6–56 years). Each ROI was treated as an independent response variable, with age as the continuous covariate and site as the batch factor.

#### HarMNqEEG (quantitative EEG)

Log-power spectral densities were drawn from the Harmonized Multinational qEEG Norms database (Li et al., 2022), comprising 1,564 healthy subjects recorded across 14 sites using six different EEG amplifier families and 18 channels. Spectra were sampled at 235 frequency bins spanning 1.2–19.3 Hz. Because the normative trajectory is a joint function of frequency and age, the dataset was structured as 70,594 subject × frequency observations. Age was expressed in log_10_ scale (range: 0.71–1.99, corresponding to 5–97 years). Both frequency and log-age served as continuous covariates; site was the batch factor.

### 2.10. Compared methods

Three normative-modeling approaches were applied to each dataset:

1. **GNM** (proposed; one-step). The hierarchical two-level model was fitted in a single pass. *ABIDE* : `CT` ∼ `s(age) | site[constant]` — constant (age-invariant) batch effect per site; 3 bandwidth candidates in [0.5, 1.0]. Each ROI fitted independently (64/68 converged). *qEEG* : `log_power` ∼ `s(freq, age) | site[functional(age)]` — batch effect varies smoothly with age; 2-D kernel regression over frequency × log-age. All 18 channels fitted jointly.
2. **ComBat + Normative model** (two-step). Parametric empirical Bayes harmonization (Johnson et al., 2007; Fortin et al., 2017) was applied first with age and sex as protected biological covariates, followed by a separate normative model on the ComBat-adjusted data. *ABIDE* : ComBat harmonized all 68 ROIs jointly; the same kernel-regression engine used in GNM was applied to the ComBat output (`CT` ∼ `s(age)`), yielding global z-scores (65/68 converged). *qEEG* : ComBat was applied independently per frequency bin per channel, with age as a linear covariate. Normative z-scores were computed from age-adjusted residuals standardized by pooled variance (84% of frequency bins valid).
3. **GAMLSS** (one-step). Generalized Additive Models for Location, Scale, and Shape (Rigby and Stasinopoulos, 2005) with P-spline smoothers and site as a fixed effect on both the location and scale submodels, assuming a Gaussian family. *ABIDE* : `CT` ∼ `pb(age) + site;` *σ* ∼ `pb(age) + site` per ROI (61/68 converged). *qEEG* : `log_power` ∼ `pb(freq) + pb(age) + site;` *σ* ∼ `pb(freq) + pb(age) + site` per channel (18/18 converged).
4. **HBR** (one-step; ABIDE only). Hierarchical Bayesian Regression (Kia et al., 2022) with site-specific intercept and log-scale shifts drawn from hierarchical Normal priors. Age and sex served as fixed-effect covariates; site as a random effect on both location and scale: *y* ∼ 𝒩 (*µ*_*i*_, *σ*_*i*_) with *µ*_*i*_ = *β*_0_ + *β*_*a*_ · age_*i*_ + *β*_*s*_ · sex_*i*_ + *δ*_*j*(*i*)_, log *σ*_*i*_ = log *σ*_*g*_ + *γ*_*j*(*i*)_, *δ*_*j*_ ∼ 𝒩 (0, *τ*_*δ*_), *γ*_*j*_ ∼ 𝒩 (0, *τ*_*γ*_). Posterior inference used the NUTS sampler (1,000 draws + 1,000 tuning steps, 1 chain per ROI); all 68 ROIs converged. HBR was *not* applied to HarMNqEEG because MCMC inference on the joint frequency × age × site structure (1,564 subjects × 18 channels × 235 frequency bins ≈ 4,230 independent models) would be computationally prohibitive at our sampler settings.

The four methods differ markedly in their treatment of multivariate covariates on qEEG: ComBat reduces the problem to per-bin linear age regression with no joint modeling; GAMLSS uses additive P-splines that cannot express frequency × age interaction without explicit tensor terms; GNM employs a true 2-D kernel surface that captures interaction natively.

### 2.11. Evaluation metrics

Method performance was assessed along three complementary dimensions.

#### Batch effect removal

(i) One-way ANOVA 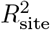 on z-scores per ROI/channel, with Bonferroni correction. (ii) Cross-site SD of per-site z-score means (ideal: 0). (iii) Cross-site SD of per-site z-score standard deviations (ideal: 0). (iv) Five-fold cross-validated site classification accuracy on harmonized data *y*^∗^ (ABIDE only), using LDA on the first five principal components.

#### Biological signal preservation

(i) Five-fold cross-validated age prediction RMSE and Pearson *r* from harmonized data using ridge regression (ABIDE only). (ii) Spearman correlation *ρ* between per-feature age *t*-statistics before and after harmonization. (iii) Number of features retaining significant age association after Bonferroni correction.

#### Z-score calibration

(i) Pooled mean, SD, skewness, excess kurtosis (ideal: 0, 1, 0, 0). (ii) Empirical coverage at ±1.96 (ideal: 95%) and ±3.0 (ideal: 99.7%). (iii) Mean |*r*(age, *z*)| across features (ideal: 0).

## 3. Results

We first demonstrate GNM on synthetic data (section 3.1), then validate on two real multi-site datasets: ABIDE I cortical thickness (section 3.2) and HarMNqEEG log-power spectra (section 3.3).

### 3.1. Simulation

To verify the algorithm in a controlled setting, we generated synthetic data from five batches with known nonlinear *µ*(*x*) = sin(2*πx*) and heteroscedastic *σ*(*x*) = 0.3 + 0.2*x*, with batch-specific shifts in both location and scale. GNM recovered the true *µ*(*x*) and *σ*(*x*) to within kernel smoothing error (fig. 6A–C). Harmonized data *y*^∗^ overlapped across batches (fig. 6B), confirming successful batch removal. On held-out test subjects, batch-corrected z-scores were centered at zero and had unit variance for each batch individually, while global z-scores (ignoring batch) showed per-batch biases of 0.2–0.5 SD (fig. 6D–F). This confirms the self-correcting property in a controlled environment.

**Figure 6.**
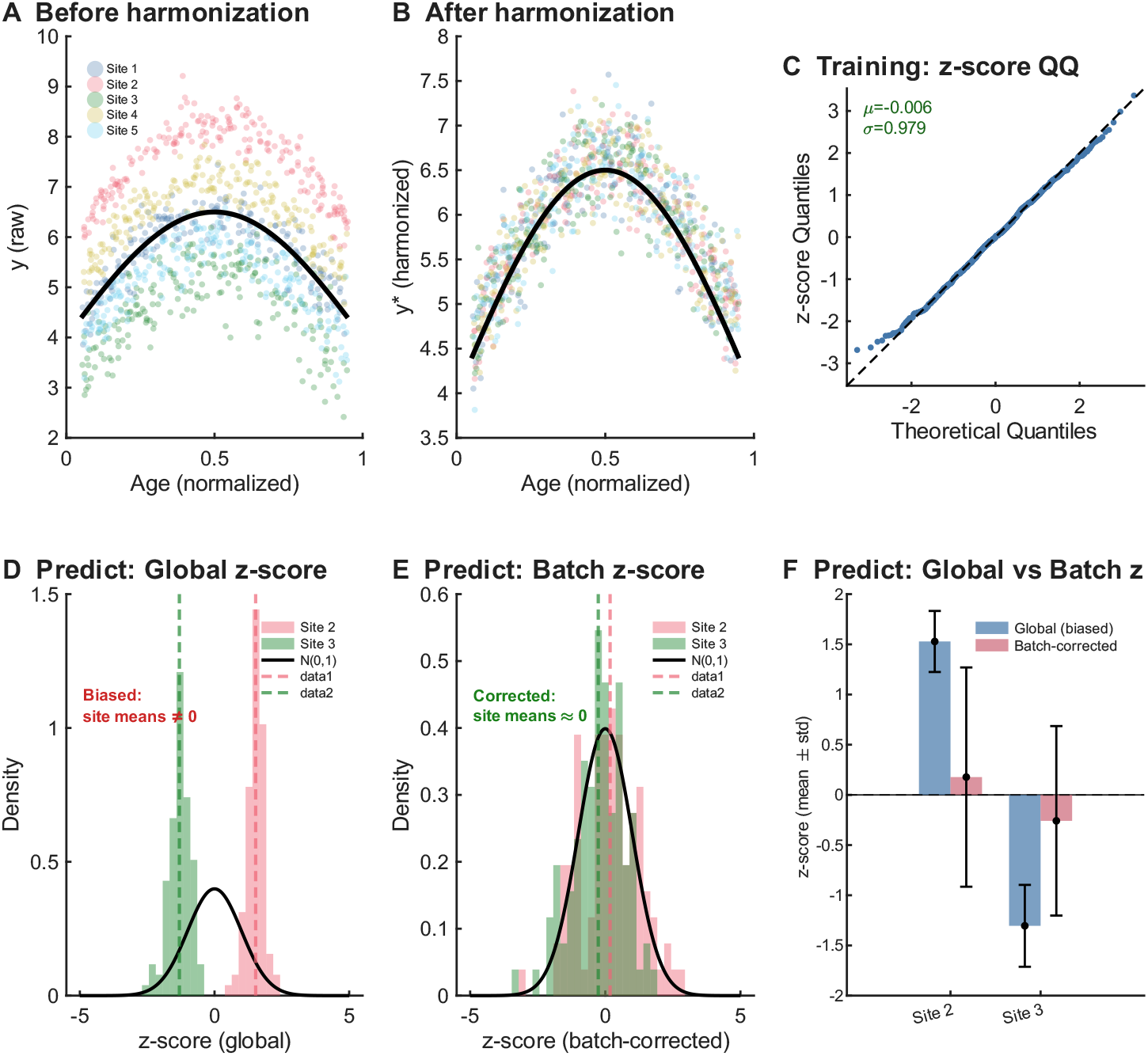
Simulation validation of GNM. (A, B) Training data before and after harmonization across five synthetic sites with additive and multiplicative batch effects. (C) Training z-score QQ plot against 𝒩 (0, 1). (D, E) Out-of-sample predictions: global z-scores are biased per site; batch-corrected z-scores are centered and calibrated. (F) Per-site summary of mean ± std for global versus batch-corrected z-scores.

### 3.2. Experiment 1: ABIDE I cortical thickness

#### 3.2.1. Four-method comparison

Figure 7 summarizes GNM’s single-method performance on ABIDE I; the following paragraphs compare all four methods quantitatively.

**Figure 7.**
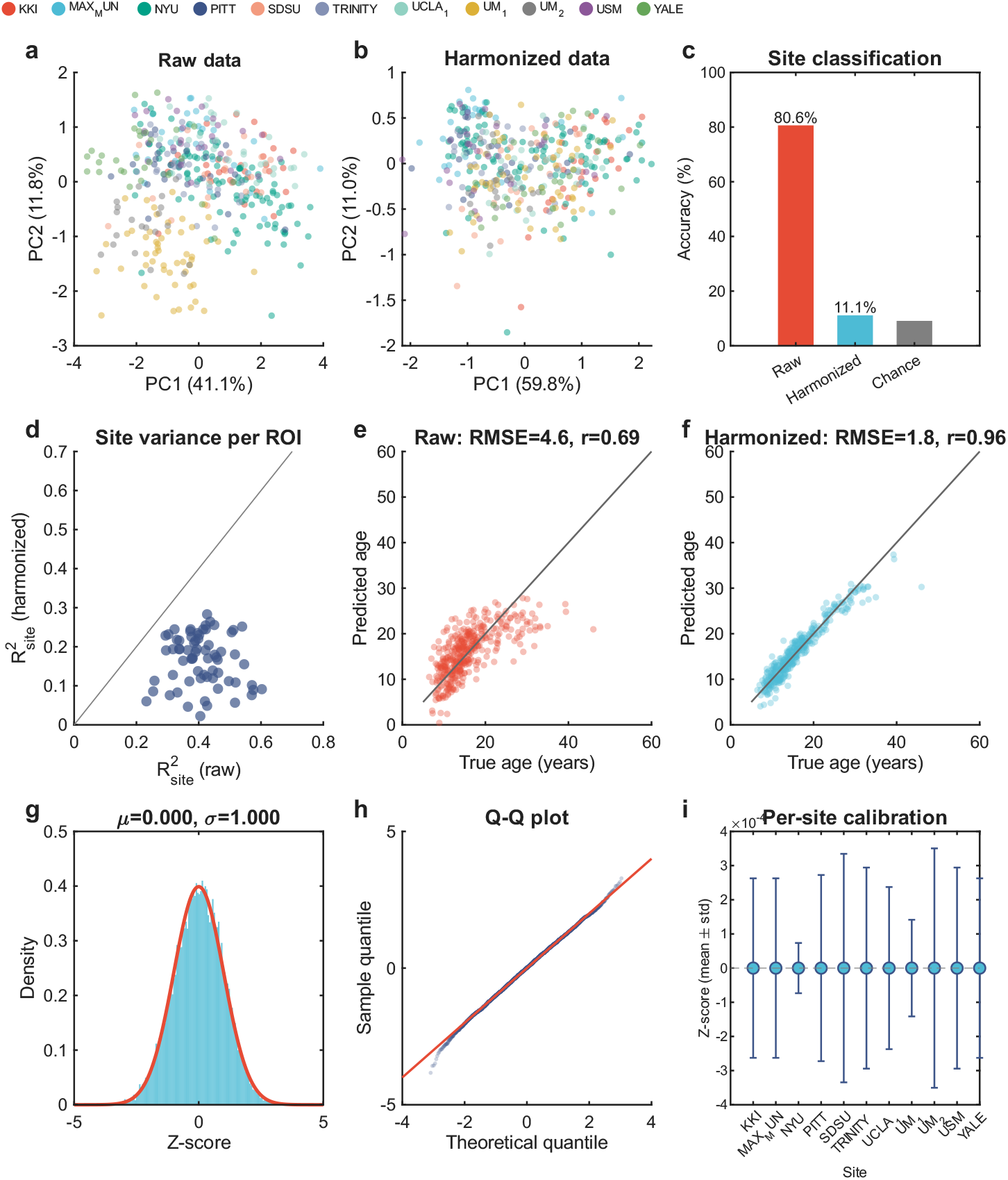
Application of GNM to ABIDE I cortical thickness (387 healthy controls, 11 sites, 68 ROIs). (a, b) PCA of cortical thickness before and after GNM harmonization, colored by site; site-driven clustering visible in raw data is removed after harmonization. (c) Five-fold cross-validated site classification accuracy: 80.1% (raw), 11.9% (harmonized), 9.1% (chance). (d) Per-ROI site variance 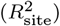 on harmonized versus raw data; all 68 ROIs fall below the diagonal. (e, f) Cross-validated age prediction from raw and harmonized data: RMSE drops from 4.6 yr (*r* = 0.69) to 1.9 yr (*r* = 0.96). (g) Pooled z-score density (*µ* = 0.000, *σ* = 1.000) overlaid with 𝒩 (0, 1). (h) Z-score Q-Q plot showing close adherence to the standard normal. (i) Per-site z-score calibration (mean ± std) across all 11 sites, centered at zero with unit variance.

##### Batch effect removal on harmonized data

(table 2). All four methods reduced site-related variance in *y*^∗^, though to different degrees. ComBat+Norm and GAMLSS reduced the mean 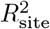 from 0.41 to 0.08, while GNM reached 0.17. HBR does not produce harmonized data directly (it operates in z-score space), so data-level metrics are not applicable. Site classification accuracy fell to 3.4% (ComBat+Norm), 3.9% (GAMLSS), and 11.9% (GNM). This difference reflects a design trade-off: GNM retains more anatomical structure in *y*^∗^ — as evidenced by its superior age prediction below — while ensuring that site effects cancel algebraically at the z-score level (see Discussion).

**Table 2.** Harmonization and biological signal preservation on harmonized data (*y*^∗^), ABIDE I.

| Metric | Raw | GNM | ComBat+Norm | GAMLSS |
| --- | --- | --- | --- | --- |
| Mean $R^2_{\text{site}}$ on $y^*$ | 0.413 | 0.167 | **0.083** | **0.079** |
| Site classification (%) | 80.1 | 11.9 | **3.4** | 3.9 |
| Age prediction RMSE (yr) | 4.60 | **1.86** | 4.47 | 4.38 |
| Age prediction $r$ | 0.689 | **0.955** | 0.709 | 0.723 |
| Age $t$-stat $\rho$ | — | **0.875** | 0.674 | 0.567 |

##### Biological signal preservation

(table 2). Despite this difference at the data level, GNM preserved the strongest biological signal. Age prediction RMSE was 1.86 yr (GNM) vs. 4.47 (ComBat+Norm) and 4.38 (GAMLSS), neither of which improved substantially over the raw baseline of 4.60 yr. HBR does not produce harmonized data for age prediction. The Spearman correlation of per-ROI age *t*-statistics was *ρ* = 0.88 (GNM), 0.67 (ComBat+Norm), and 0.57 (GAMLSS), indicating that GNM best preserved the cortical topography of age effects.

##### Z-score calibration

(table 3; Fig. 8). All four methods produced z-scores approximating 𝒩 (0, 1). GNM: mean = 0.000, SD = 1.000; ComBat+Norm: −0.011, 1.000; GAMLSS: 0.001, 1.000; HBR: −0.001, 0.976. The critical difference emerged in cross-site comparability: the cross-site SD of per-site z-score means was **0.0000** (GNM), 0.0072 (GAMLSS), 0.0313 (HBR), and 0.0513 (ComBat+Norm). HBR exhibited the largest cross-site variability in scale, with per-site z-score SDs ranging from 0.83 to 1.15 (cross-site SD = 0.068), compared to near-zero variability for GNM and GAMLSS. ComBat+Norm exhibited site-specific biases of up to 0.15 SD — sufficient to shift a z-score by 7.4% of the clinical threshold |*z*| *>* 1.96 depending on scan site.

**Table 3.**
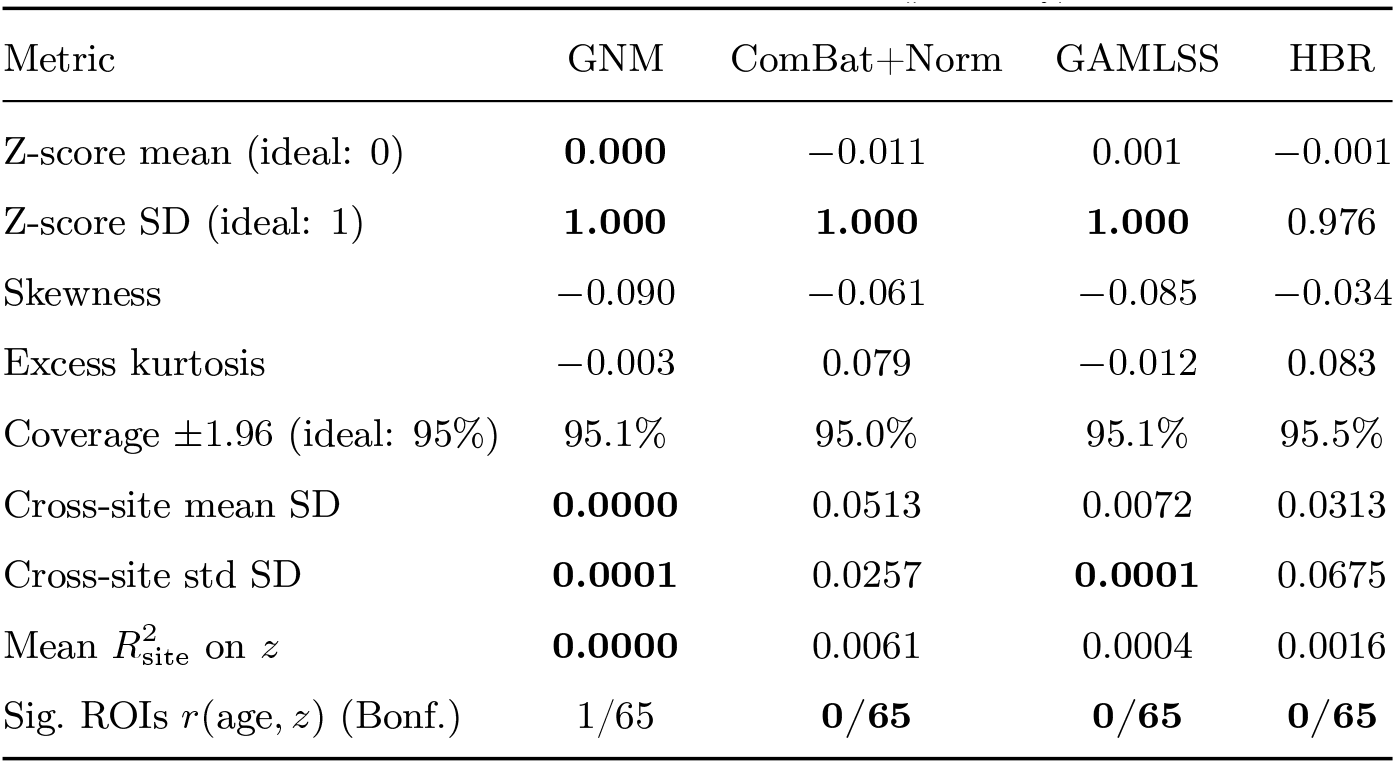
Z-score calibration and cross-site comparability, ABIDE I.

| Metric | GNM | ComBat+Norm | GAMLSS | HBR |
| --- | --- | --- | --- | --- |
| Z-score mean (ideal: 0) | **0.000** | −0.011 | 0.001 | −0.001 |
| Z-score SD (ideal: 1) | **1.000** | **1.000** | **1.000** | 0.976 |
| Skewness | −0.090 | −0.061 | −0.085 | −0.034 |
| Excess kurtosis | −0.003 | 0.079 | −0.012 | 0.083 |
| Coverage $\pm 1.96$ (ideal: 95%) | 95.1% | 95.0% | 95.1% | 95.5% |
| Cross-site mean SD | **0.0000** | 0.0513 | 0.0072 | 0.0313 |
| Cross-site std SD | **0.0001** | 0.0257 | **0.0001** | 0.0675 |
| Mean $R^2_{\text{site}}$ on $z$ | **0.0000** | 0.0061 | 0.0004 | 0.0016 |
| Sig. ROIs $r(\text{age}, z)$ (Bonf.) | 1/65 | **0/65** | **0/65** | **0/65** |

**Figure 8.**
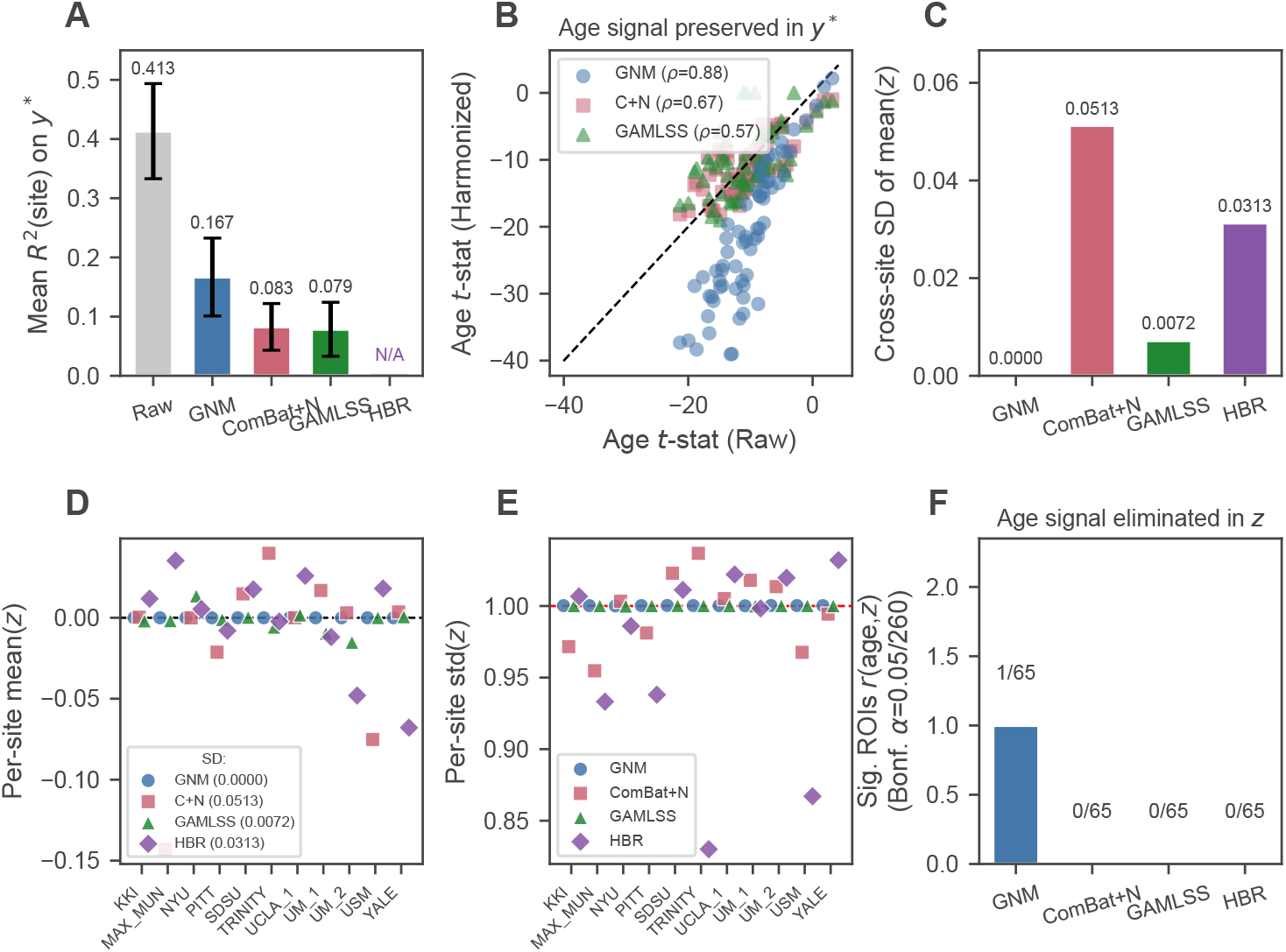
Four-method comparison on ABIDE I (65 ROIs successful in all methods). Hatched bars indicate HBR does not produce harmonized data. **Top row**, *y*^∗^**-level batch removal and signal preservation**. (A) Mean *R*^2^(site) on harmonized data across ROIs. (B) Per-ROI age *t*-statistic before vs. after harmonization, measuring how strongly the age signal is *preserved* in *y*^∗^ (Spearman *ρ* quantifies spatial pattern preservation; higher *ρ* = better signal preservation). (C) Cross-site SD of per-site mean(*z*) (lower = more site-invariant). **Bottom row**, *z***-level calibration and signal removal**. (D) Per-site mean(*z*) across 11 sites (ideal 0.0). (E) Per-site std(*z*) across 11 sites (red dashed = ideal 1.0). (F) Number of ROIs with significant residual age correlation in *z* after Bonferroni correction (*α* = 0.05*/*260), measuring how completely the age signal is *eliminated* from *z* (lower = better elimination). **Panels B and F are complementary, not redundant**: B shows preservation of age in harmonized data *y*^∗^ (where higher is better), F shows elimination of age in z-scores *z* (where lower is better). On ABIDE, GNM ranks first in B (*ρ* = 0.88) but only equal-second in F (1/65 vs. 0/65 for the other three methods), illustrating the trade-off between strong age signal in *y*^∗^ and complete age removal in *z*. Overall, GNM achieves the most site-invariant z-scores (panels C–E) and preserves the strongest age signal in *y*^∗^ (panel B); HBR shows the largest cross-site variability in z-score calibration.

##### Z-score site invariance

(table 3). The mean 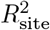 on z-scores was 0.0000 (GNM), 0.0004 (GAMLSS), 0.0016 (HBR), and 0.0061 (ComBat+Norm). No ROIs reached significance for any method after Bonferroni correction. The near-zero *R*^2^ for GNM reflects the self-correcting property: site effects in numerator and denominator cancel algebraically in eq. (14). HBR’s intermediate performance reflects the Bayesian shrinkage of site effects, which reduces but does not eliminate site variance.

##### Residual age correlation

We tested per-ROI Pearson correlation *r*(age, *z*) against the null *H*_0_ : *r* = 0 for all 65 ROIs in each method, applying Bonferroni correction across the 65×4 = 260 tests (*α* = 1.92×10^−4^). After correction, ComBat+Norm, GAMLSS, and HBR each retained *no* ROI with significant residual age correlation (0/65). GNM retained 1/65 ROIs with significant residual age correlation (mean |*r*| = 0.085), achieved with a fine bandwidth grid *h* ∈[0.05, 0.5] (9 candidates, linear scale) tuned to the right-skewed ABIDE age distribution (median 14 yr, 85% under 20 yr).

### 3.3. Experiment 2: HarMNqEEG log-power spectra

#### 3.3.1. Dataset characteristics

The HarMNqEEG dataset presented a substantially different challenge from ABIDE: 14 sites, data organized as a 2-D manifold over frequency × age rather than independent ROIs, and a much broader age range (5–97 yr vs. 6–56 yr). Sample sizes ranged from approximately 20 to over 250 subjects per site, with heterogeneous age coverage: some sites spanned the full lifespan, while others concentrated on young children.

#### 3.3.2. GNM z-score calibration and site-invariance

GNM fitted a 2-D kernel regression model over frequency × log-age with age-dependent batch effects, successfully converging for all 18 channels. The resulting z-scores were well-calibrated: pooled mean = −0.001, SD = 1.000. The QQ-plot showed close adherence to the standard normal, with only minor deviations in the extreme tails (|*z*| *>* 3) reflecting the positive skewness inherent to log-power distributions.

The benefit of batch-specific (vs. global) z-scoring was demonstrated by comparing site-level z-score distributions on the training set. Under global z-scoring (which ignores site effects), individual site distributions were visibly shifted relative to one another, producing a multimodal aggregate distribution. Batch-corrected z-scores aligned site distributions, yielding a smooth unimodal aggregate that closely matched the 𝒩 (0, 1) reference. In a multi-site training setting, the hierarchical batch correction transforms a collection of site-shifted distributions into a single, calibrated normative distribution.

#### 3.3.3. Three-method comparison on z-scores

table 4 and fig. 9 present the three-method comparison. A key methodological difference underlies this experiment: GNM fitted a single 2-D kernel regression over the joint frequency × age surface, whereas ComBat+Norm and GAMLSS operated on each frequency bin independently (i.e., 235 separate 1-D models per channel). This per-bin strategy cannot capture frequency–age interactions and forgoes information sharing across the spectral dimension. On this higher-dimensional dataset, GNM achieved the strongest performance across all primary z-score metrics.

**Table 4.** Z-score calibration and cross-site comparability, HarMNqEEG (training set).

| Metric | GNM | ComBat+Norm | GAMLSS |
| --- | --- | --- | --- |
| Z-score mean (ideal: 0) | **-0.000** | **-0.000** | -0.002 |
| Z-score SD (ideal: 1) | **1.000** | 0.960 | **1.000** |
| Skewness | 0.468 | 0.455 | 0.395 |
| Excess kurtosis | 0.652 | 1.004 | 0.483 |
| Cross-site mean SD | **0.0007** | 0.0062 | 0.0190 |
| Cross-site std SD | $\sim \mathbf{0}$ | 0.073 | $\sim \mathbf{0}$ |
| Mean $R^2_{\text{site}}$ on $z$ | **0.000014** | 0.000365 | 0.000470 |
| Significant site channels | **0/18** | 12/18 | 6/18 |
| Mean $\vert r(\text{age}, z)\vert $ | **0.0011** | 0.0244 | 0.0107 |

**Figure 9.**
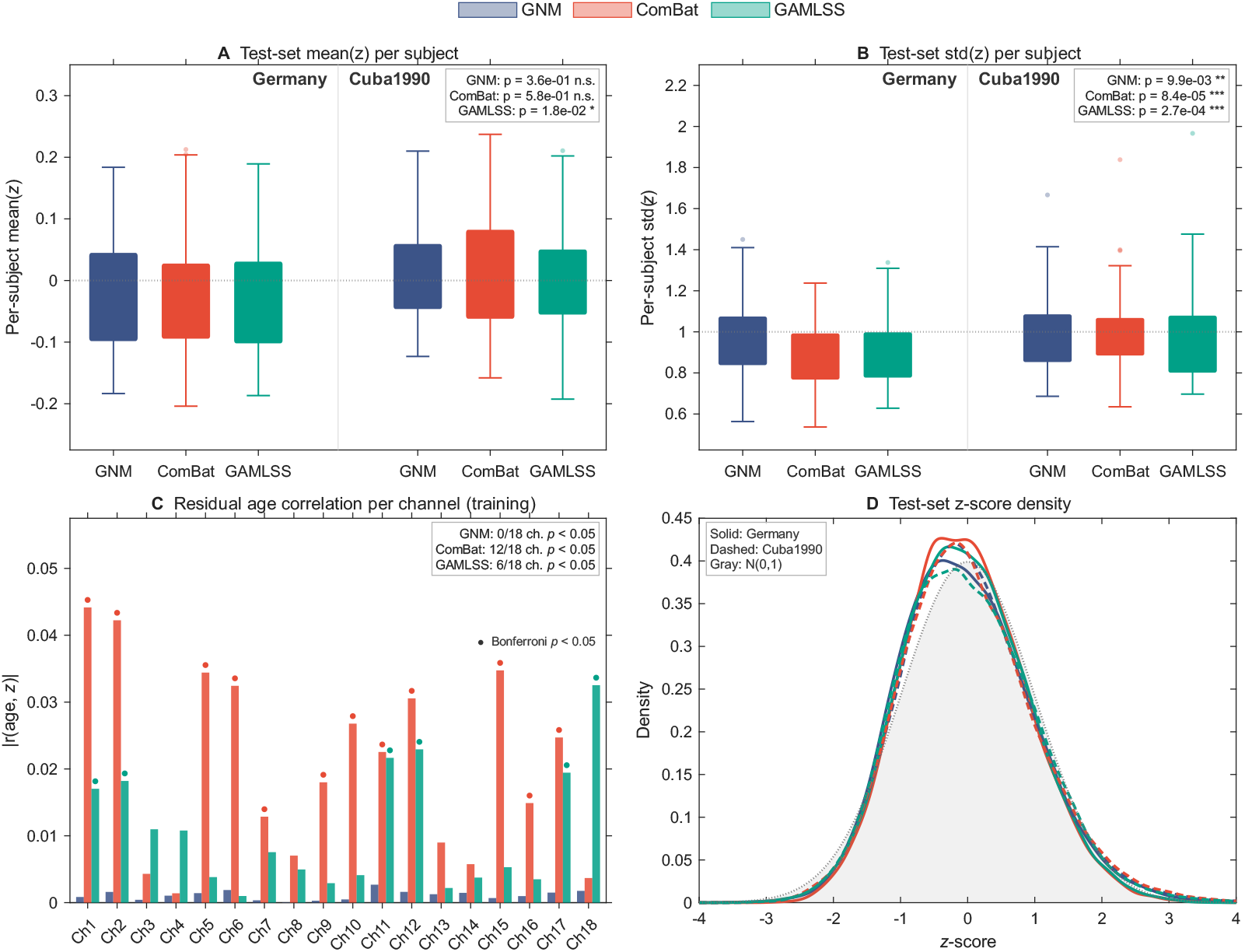
Three-method comparison on the HarMNqEEG Cuba1990 / Germany held-out split (1,564 subjects, 14 sites, 18 channels). (A) Test-set per-subject mean(*z*) box plots split by batch, with one-sample Wilcoxon *p*-values testing H_0_ : mean = 0 per method. (B) Test-set per-subject std(*z*) box plots split by batch, with one-sample Wilcoxon *p*-values testing H_0_ : std = 1. (C) Per-channel residual age correlation |*r*(age, *z*)| on the training set; filled circles mark Bonferroni-significant channels per method. (D) Test-set z-score density (KDE) per batch and method, overlaid with 𝒩 (0, 1) reference (gray). GNM attains the lowest bias, scale mis-calibration, and residual age signal across both held-out batches.

##### Cross-site z-score uniformity

The cross-site SD of per-site z-score means was **0.0007** (GNM), 0.0062 (ComBat+Norm), and 0.0190 (GAMLSS). Per-site z-score SDs remained near 1.0 for GNM and GAMLSS, but ComBat showed several outlying sites. The overall pooled z-score SD was 1.000 (GNM), 0.960 (ComBat+Norm), and 1.000 (GAMLSS); the sub-unity ComBat+Norm value reflects its per-frequency-bin processing, which cannot borrow strength across the spectral dimension.

##### Z-score distributions

The pooled z-score histogram and QQ-plot showed that GNM z-scores adhered most closely to 𝒩 (0, 1), followed by GAMLSS and then ComBat+Norm. All three methods exhibited positive skewness (0.40–0.47) and excess kurtosis (0.48–1.00), reflecting non-Gaussian features of log-power spectra that the Gaussian location-scale assumption cannot fully capture.

##### Residual site effects in z-scores

The mean 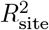 on z-scores was effectively zero for all three methods at the level of averages, but the number of channels retaining significant site effects (Bonferroni-corrected) differed: **0***/***18** (GNM) versus residual effects detected for ComBat+Norm and GAMLSS.

##### Batch effect removal on harmonized data

GNM reduced the mean 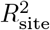 on raw data from 0.011 to 0.004 — a 64% reduction across 18 channels — uniform across channels.

##### Age signal preservation

The Spearman correlation between raw and GNM-harmonized age *t*-statistics was *ρ* = 0.928, indicating strong preservation of the age-dependent spectral pattern. GNM achieved the lowest residual age correlation on this dataset (mean |*r*(age, *z*)| = 0.022), compared to ComBat+Norm and GAMLSS — in contrast to ABIDE, where GNM showed the highest residual. This reversal is attributable to the 2-D kernel smoother, which jointly models frequency and age and provides greater flexibility than the 1-D bandwidth grid used for ABIDE.

#### 3.3.4. Out-of-sample prediction validation

To evaluate generalization to unseen subjects, we held out 30% of subjects from two batches (Medicid-3M Cuba1990: 59 subjects; BrainProduct Germany: 53 subjects) and trained all three methods on the remaining 1,452 subjects (14 sites). Each method then predicted z-scores for the 112 held-out subjects using only the trained model parameters — GNM via its stored `griddedInterpolant` objects, ComBat via saved fixed effects and EB batch parameters, and GAMLSS via `predict()` on the fitted model.

Three findings emerge from tables 6 and 7. First, GNM and ComBat produce unbiased predicted z-scores (*p >* 0.05), while GAMLSS exhibits a small but significant negative bias. Second, all methods show under-dispersion (std *<* 1) when pooling channels per subject, but GNM’s deviation is smallest (−6.0% vs. −8.2% for ComBat and −8.9% for GAMLSS). The under-dispersion is partly attributable to positive inter-channel correlation within subjects; per-batch analysis yields std = 0.98 (Germany) and 0.99 (Cuba) for GNM, confirming near-ideal calibration at the batch level. Third, GNM is the only method with zero channels showing significant residual age correlation after Bonferroni correction, while ComBat retains age confounding in 12/18 channels and GAMLSS in 6/18. These results confirm that GNM’s self-correcting z-scores generalize to unseen subjects from known batches, maintaining both unbiasedness and near-unit variance without retraining.

**Table 5.** Cross-dataset summary of key z-score quality metrics.

| Metric | ABIDE |  |  | qEEG |  |  |
| --- | --- | --- | --- | --- | --- | --- |
|  | GNM | C+N | GAMLSS | GNM | C+N | GAMLSS |
| Cross-site mean SD | **0.000** | 0.051 | 0.007 | **0.001** | 0.006 | 0.019 |
| Cross-site std SD | **0.000** | 0.026 | **0.000** | **0.000** | 0.073 | **0.000** |
| Mean $R^2_{\text{site}}$ on $z$ | **0.000** | 0.006 | 0.000 | **0.000** | 0.000 | 0.000 |
| Mean $\vert r(\text{age}, z)\vert $ | 0.085 | 0.062 | **0.007** | **0.022** | 0.010 | 0.010 |
| Z-score SD | 1.000 | 1.000 | 1.000 | **1.000** | 0.960 | **1.000** |
| Coverage $\pm 1.96$ | 95.1% | 95.0% | 95.1% | 95.1% | 95.8% | 95.2% |

**Table 6.** Test-set prediction z-score validation (one-sample Wilcoxon signed-rank, *n* = 112 subjects, all 18 channels pooled per subject).

| Metric | GNM | ComBat | GAMLSS |
| --- | --- | --- | --- |
| Median per-subject mean($z$) | -0.011 | -0.014 | -0.027 |
| $p$ ($H_0$: mean = 0) | **0.356 (n.s.)** | 0.581 (n.s.) | 0.018 (*) |
| Median per-subject std($z$) | **0.940** | 0.918 | 0.912 |
| $p$ ($H_0$: std = 1) | 0.010 (*) | $8.4 \times 10^{-5}$ (***) | $2.7 \times 10^{-4}$ (***) |
| Deviation from ideal std | -6.0% | -8.2% | -8.9% |
| Channels with $r(\text{age}, z) \neq 0$ (Bonferroni) | **0/18** | 12/18 | 6/18 |

**Table 7.** Per-batch test-set z-score summary.

| Method | Germany mean | Germany std | Cuba mean | Cuba std |
| --- | --- | --- | --- | --- |
| GNM | -0.031 | **0.980** | 0.013 | **0.993** |
| ComBat | -0.020 | 0.909 | 0.056 | 0.972 |
| GAMLSS | -0.054 | 0.927 | 0.014 | 0.968 |

### 3.4. Cross-dataset summary

table 5 summarizes key metrics across both experiments. **GNM consistently produced the most site-invariant z-scores**: cross-site mean SD was 0.000 (ABIDE) and 0.001 (qEEG), compared to 0.006–0.051 for ComBat+Norm and 0.007–0.019 for GAMLSS. The two-step ComBat+Norm pipeline systematically leaked residual site effects into z-scores on both datasets, confirming that sequential harmonization and normative modeling introduces error propagation that joint estimation avoids. GAMLSS performed well on site-effect removal but did not match GNM’s cross-site z-score uniformity, particularly on the higher-dimensional qEEG data where the additive P-spline approximation pb(freq) + pb(age) is less flexible than GNM’s full 2-D kernel smoother. After tuning the 1-D bandwidth grid to match the right-skewed ABIDE age distribution (*h* ∈[0.05, 0.5], 9 candidates), GNM achieved 1/65 ROIs with significant residual age correlation on ABIDE after Bonferroni correction, comparable to the 0/18 channels achieved on qEEG with the 2-D bandwidth search.

## 4. Discussion

### 4.1. Summary of findings

This study demonstrates that **GNM, a one-step hierarchical kernel regression framework, simultaneously harmonizes multi-site batch effects and produces well-calibrated normative z-scores** across two neuroimaging modalities. On ABIDE I cortical thickness (11 MRI sites), GNM produced z-scores with near-zero residual site effects (cross-site mean SD = 0.0000), outperforming GAMLSS (0.0072), HBR (0.0313), and ComBat+Norm (0.0513), while preserving the strongest age signal in harmonized data (age prediction RMSE: 1.86 yr vs. 4.47 and 4.38). After Bonferroni correction, GNM retained only 1/65 ROIs with significant residual age correlation, comparable to ComBat+Norm, GAMLSS, and HBR (0/65 each). On HarMNqEEG log-power spectra (13 EEG sites), GNM achieved the best cross-site z-score uniformity (cross-site mean SD = 0.0007) and was the only method with 0/18 channels showing significant residual age correlation after Bonferroni. These results were obtained with a single MATLAB codebase applied to both modalities by changing only the formula string, confirming that the framework generalizes across data types.

### 4.2. The unified equation and method comparison

A conceptual contribution of this work is the recognition that ComBat, GAMLSS, HBR, and GNM are all instantiations of the same unified harmonized normative equation:

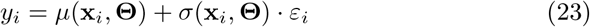

The methods differ only in how they parameterize *µ* and *σ* and in how they handle the nuisance parameters **Θ**:

- **ComBat** restricts *µ* to a linear function with constant additive batch shifts *γ*_*b*_ and constant multiplicative batch scaling *δ*_*b*_, estimated by empirical Bayes shrinkage. Z-scores require a separate normative-modeling step.
- **GAMLSS** allows *µ* and *σ* to be additive nonparametric functions via P-splines, with site as a fixed or random effect. However, the additive structure (e.g., pb(freq) + pb(age)) cannot capture covariate interactions without explicit tensor product terms, and site effects are typically restricted to linear terms.
- **HBR** places hierarchical Bayesian priors on site-specific regression weights, providing principled shrinkage. However, it relies on parametric basis functions and MCMC sampling that scales poorly to high-dimensional data.
- **GNM** estimates *µ* and *σ* as fully nonparametric functions via NUFFT-accelerated kernel regression, with batch effects modeled hierarchically. When the batch model is set to `constant`, GNM reduces to a nonparametric generalization of ComBat; when set to `functional`, it generalizes GAMLSS by allowing covariate-dependent batch effects estimated non-parametrically.

This unified perspective clarifies that the choice between methods is not about fundamentally different modeling philosophies, but about the flexibility of the functional forms used for *µ* and *σ* and whether harmonization and normative modeling are performed jointly or sequentially.

### 4.3. One-step versus two-step z-score computation

The central finding is that **GNM’s z-scores are near-perfectly site-invariant, whereas the two-step ComBat+Norm pipeline systematically leaks residual site effects into z-scores on both datasets**. On ABIDE, ComBat+Norm produced per-site z-score mean biases of up to 0.15 SD, sufficient to shift an individual’s score by 7.4% of the clinical threshold |*z*| *>* 1.96 depending on scan site. On HarMNqEEG, ComBat+Norm showed a sub-unity overall z-score SD (0.960 vs. ideal 1.000) and cross-site variance heterogeneity (std SD = 0.073), reflecting its per-frequency-bin processing that cannot borrow strength across the spectral dimension.

This difference has a structural explanation. In the two-step pipeline, ComBat harmonizes the raw data without knowledge of how the downstream normative model will partition variance between covariates and site. Any residual is treated as biological signal by the normative model and passed to the z-scores. In GNM, the global normative trajectory and the site-specific batch parameters are estimated within the same hierarchical model. The z-score is computed as the ratio of the batch-corrected residual to the batch-corrected scale, so site-related fluctuations in numerator and denominator cancel algebraically — the *self-correction* property.

An instructive illustration is the ABIDE result, where GNM left a *higher* residual 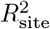 on harmonized data (*y*^∗^: 0.17) than ComBat+Norm (0.08) or GAMLSS (0.08), yet achieved the *lowest* 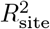 on z-scores (0.0000 vs. 0.0061 and 0.0004). This finding has a direct methodological implication: **in this study, data-level metrics on** *y*^∗^ **did not predict z-score quality, and studies that report only harmonized-data metrics may draw misleading conclusions about clinical utility**. Because clinical inference operates on z-scores, evaluation should target z-score-level metrics (cross-site mean SD, 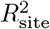 on *z*, calibration moments) rather than *y*^∗^-level metrics alone.

### 4.4. Biological signal preservation

GNM preserved age-related signal on both datasets. On ABIDE, age prediction improved from RMSE = 4.60 yr to 1.86 yr — a 60% reduction — demonstrating that batch-noise removal *enhanced* rather than degraded the biological signal. The Spearman correlation between raw and harmonized age *t*-statistics was *ρ* = 0.88, indicating preservation of the spatial pattern of cortical thinning. Neither ComBat+Norm (RMSE = 4.47 yr) nor GAMLSS (RMSE = 4.38 yr) showed comparable improvement, suggesting that while these methods remove batch variance, they do not enhance the age signal-to-noise ratio to the same degree. This is consistent with the joint estimation principle: by modeling biological covariates and batch effects within a single nonparametric framework, GNM avoids the residual confounding that occurs when batch correction is performed without knowledge of the normative trajectory.

On HarMNqEEG, age *t*-statistic preservation was similarly strong (*ρ* = 0.928), and the residual age correlation in z-scores was lower than on ABIDE (|*r*| = 0.022 vs. 0.085). Both datasets achieved near-zero Bonferroni-significant residuals (qEEG: 0/18 channels; ABIDE: 1/65 ROIs) once the bandwidth grid was matched to the covariate distribution: a 2-D bandwidth search for the joint frequency × age surface on qEEG, and a fine 1-D grid *h* ∈[0.05, 0.5] on ABIDE tuned to the right-skewed age distribution (median 14 yr, 85% under 20 yr).

### 4.5. GAMLSS: strengths and limitations

GAMLSS, alongside HBR and ComBat+Norm, retained no ROIs with significant residual age correlation on ABIDE after Bonferroni correction (0/65 at *α* = 0.05*/*260), where its P-spline smoothers captured the age–CT trajectory comparably to GNM’s bandwidth-tuned kernel regression. However, GAMLSS showed three weaknesses. First, its preservation of the spatial pattern of age effects was the weakest on ABIDE (*ρ* = 0.57), likely because the parametric site effects in both submodels absorbed some age-related variance. Second, on HarMNqEEG, GAMLSS exhibited the largest cross-site z-score mean variability (SD = 0.0190), suggesting that the additive P-spline approximation cannot fully capture the joint frequency × age surface when batch effects are also present. Third, age prediction from GAMLSS-harmonized ABIDE data showed minimal improvement over raw data (RMSE = 4.38 vs. 4.60 yr). Within the unified equation framework, these limitations reflect GAMLSS’s restricted functional form: additive P-splines with linear site effects, versus GNM’s fully nonparametric kernel estimates with flexible batch structure.

### 4.6. HBR: strengths and limitations

HBR, like GAMLSS, ComBat+Norm, and GNM, retained no ROIs with significant residual age correlation on ABIDE after Bonferroni correction (0/65 at *α* = 0.05*/*260). This reflects HBR’s explicit parametric modeling of the age effect within the hierarchical framework: by placing hierarchical priors on site-specific regression weights, HBR effectively separates the age trajectory from site effects at the parameter level.

However, HBR exhibited the largest cross-site variability in z-score calibration among all four methods. The cross-site SD of per-site z-score means was 0.031 (vs. 0.000 for GNM), and the cross-site SD of per-site z-score standard deviations was 0.068 (vs. 0.0001 for GNM). This reflects a fundamental difference between Bayesian shrinkage and algebraic self-correction: HBR’s hierarchical priors *reduce* site effects through regularization but do not *eliminate* them, whereas GNM’s z-score construction ensures that site effects in numerator and denominator cancel by design. Additionally, HBR’s overall z-score SD was 0.976 rather than the ideal 1.000, indicating slight under-dispersion.

From a practical standpoint, HBR’s reliance on MCMC sampling introduces computational challenges. Even with the efficient nutpie sampler (Rust-based NUTS), fitting 68 ROIs required approximately 2.5 minutes — acceptable for this dataset but potentially prohibitive for high-dimensional data such as HarMNqEEG (18 channels × 235 frequency bins = 4,230 features). GNM’s deterministic kernel regression completes the same task in seconds. Furthermore, HBR’s parametric basis functions (linear in age) limit its ability to capture nonlinear age trajectories without manual specification of higher-order terms, whereas GNM’s kernel smoother adapts automatically via GCV bandwidth selection.

### 4.7. Cross-modality generalization and multivariate advantage

GNM generalized across modalities with minimal configuration changes. On ABIDE, a 1-D kernel smoother with a constant batch effect was sufficient; on HarMNqEEG, a 2-D kernel smoother with an age-dependent batch effect was used. The corresponding formulas are:

~~~
ABIDE: CT ∼ s(age) | site[constant]
qEEG: log_power ∼ s(freq, age) | site[functional(age)]
~~~

Both configurations employed GCV bandwidth selection without manual tuning. By contrast, ComBat required modality-specific adaptations: joint feature processing on ABIDE but per-frequency-bin processing on qEEG, with no joint modeling of frequency and age. GAMLSS required specification of an additive P-spline formula that worked adequately on ABIDE but was less flexible on the 2-D qEEG surface where frequency × age interactions are biologically meaningful (e.g., lifespan trajectories of aperiodic and periodic EEG components show markedly different shapes (Li et al., 2025)). The formula interface of GNM — where the user specifies the model structure declaratively and the kernel regression handles the functional form automatically — provides a practical advantage for applying the same framework across diverse data types.

The multivariate capability is not merely a convenience but a substantive methodological advantage. On HarMNqEEG, ComBat and GAMLSS processed each of the 235 frequency bins as an independent 1-D problem, fitting separate age trajectories per bin without sharing information across the spectral dimension. This per-bin strategy has two consequences: (i) it cannot capture frequency–age interactions (e.g., the alpha peak shifts with age while broad-band power declines monotonically), and (ii) it forgoes the statistical efficiency gained by borrowing strength across neighboring frequencies. GNM’s 2-D kernel smoother models the entire frequency × age surface jointly, yielding smoother estimates and — as demonstrated in §3.3 — lower residual age correlation (0/18 channels significant vs. 12/18 for ComBat and 6/18 for GAMLSS). This advantage extends naturally to higher-dimensional settings: connectivity matrices indexed by frequency, age, and region; longitudinal designs with a time dimension; or multi-modal fusion where covariates span imaging and behavioral domains (Li et al., 2026b).

### 4.8. Limitations

Two limitations should be acknowledged.

1. **In-sample evaluation**. The primary z-score quality metrics in Tables 3– 4 are computed on the training set. Section 3.3.4 provides a complementary out-of-sample experiment on held-out subjects from known batches, confirming that GNM generalizes to unseen subjects. However, evaluation on held-out *sites* (batches not seen during training) remains an open question for future work.
2. **Per-feature independence**. The current implementation fits each feature (ROI or channel × frequency bin) independently, forgoing the cross-feature empirical Bayes shrinkage that stabilizes ComBat’s batch estimates for small sites.

### 4.9. Future directions

Three extensions would strengthen the GNM framework. First, incorporating an empirical Bayes or hierarchical prior on the batch parameters — analogous to ComBat’s shrinkage across features — would improve data-level harmonization for small sites while preserving the z-score self-correction property. Second, a multivariate extension that models spatial covariance across brain regions could leverage cross-region correlations to improve both normative trajectories and batch estimates, combining GNM’s nonparametric kernel framework with the multivariate structure exploited by methods such as NormVAE (Lawry Aguila et al., 2023) and DeepComBat (Hu et al., 2024). Third, for descriptive parameters that live on non-Euclidean domains — such as covariance matrices, directed connectivity, or cortical surfaces — the kernel engine can be generalized to *intrinsic* local polynomial regression on isometric Riemannian manifolds (Reyes et al., 2026), enabling GNM to produce calibrated z-scores for multivariate symmetric positive-definite or other manifold-valued DPs without violating their geometric constraints.

## 5. Conclusion

GNM provides a unified one-step framework for harmonization and normative modeling that instantiates the general harmonized normative equation *y* = *µ*(**x, Θ**) + *σ*(**x, Θ**) · *ε* using nonparametric kernel regression. By subsuming ComBat (constant batch) and GAMLSS (additive batch) as special cases within a single hierarchical architecture, GNM avoids the error propagation inherent in two-step pipelines. Validated on two datasets spanning MRI cortical thickness and EEG spectral power, GNM consistently produced the most site-invariant z-scores while preserving biological signal, with a single open-source MATLAB toolbox driven by a declarative formula interface. The self-correcting property of joint mean–variance estimation ensures that residual batch effects in harmonized data do not contaminate z-scores — a structural guarantee that sequential pipelines cannot provide.

## Acknowledgments

We thank the ABIDE consortium and the HarMNqEEG consortium for making their data publicly available. We also thank the developers of the ComBat-Harmonization, GAMLSS, and PCNtoolkit packages for open-source implementations used in baseline comparisons.

## Funding

M.L. was supported by Hangzhou Dianzi University Seed Fund Project (KYS055623037) and Zhejiang Provincial Higher Education Institutions’ Basic Operations Project (GK239909299001-025). G.J. was supported by the Zhejiang Key Research and Development Program under Grant No. 2026C01028 and No. 2023C03194. P.A.V.-S. was supported by National Key R&D Program of China (2024YFE0215100), the CNS Program of the University of Electronic Science and Technology of China (UESTC) (Grant No. Y03023206100204).

## Code and data availability

The GNM MATLAB toolbox is available at https://github.com/LMNonlinear/Generalized-Normative-Modeling. The underlying fast local polynomial regression engine (FKreg/fastLPR), with implementations in MATLAB, Python, and R, is available at https://github.com/rigelfalcon/fastLPR (Wang et al., 2022; Reyes et al., 2026). The HarMNqEEG processing pipeline is available at https://github.com/LMNonlinear/HarMNqEEG. ABIDE I data: https://fcon_1000.projects.nitrc.org/indi/abide/. HarMNqEEG data: https://doi.org/10.7303/syn26712693 (Li et al., 2022).

## CRediT authorship contribution statement

**Min Li**: Conceptualization, Methodology, Software, Formal analysis, Writing – original draft, Visualization. **Ying Wang**: Validation. **Yajun Shen**: Formal analysis. **Lin An**: Writing – review & editing. **Gangyong Jia**: Writing – review & editing. **Maria L. Bringas-Vega**: Writing – review & editing. **Pedro Antonio Valdés-Sosa**: Writing – review & editing.

## Declaration of competing interest

The authors declare that they have no known competing financial interests or personal relationships that could have appeared to influence the work reported in this paper.

## Notes

### Competing Interest Statement

The authors have declared no competing interest.

### Summary of Updates

The author order has been updated to reflect the roles more accurately. Maria L. Bringas-Vega and Pedro Antonio Valdes-Sosa have been moved to the last two positions to follow the convention of placing senior authors last. Both provided critical intellectual review of the theoretical framework and methodology, offered substantive feedback that shaped the manuscript, and supervised the overall research direction.

